# A parametric model for estimating the timing and intensity of animal migration

**DOI:** 10.1101/2023.01.05.522924

**Authors:** Peter R. Thompson, Peter D. Harrington, Conor D. Mallory, Subhash R. Lele, Erin M. Bayne, Andrew E. Derocher, Mark A. Edwards, Mitch Campbell, Mark A. Lewis

## Abstract

Animals of many different species, trophic levels, and life history strategies migrate, and the improvement of animal tracking technology allows ecologists to collect increasing amounts of detailed data on these movements. Understanding when animals migrate is important for managing their populations, but is still difficult despite modelling advancements. We designed a model that parametrically estimates the timing of migration from animal tracking data. Our model identifies the beginning and end of migratory movements as signaled by changes in step length and turning angle distributions. To this end, we can also use the model to estimate how an animal’s movement changes when it begins migrating. We tested our model on three datasets: migratory ferruginous hawks (*Buteo regalis*) in the Great Plains and barren-ground caribou (*Rangifer tarandus groenlandicus*) in northern Canada, and non-migratory brown bears (*Ursus arctos*) from the Canadian Arctic. We estimated the beginning and end of migration in caribou and hawks to the nearest day, while confirming a lack of migratory behaviour in the brown bear population. The flexibility of our modelling framework allowed us to assess intricacies associated with each dataset: long-term stopover behaviour in ferruginous hawks and *a priori* knowledge of caribou calving areas and behaviour. In addition to estimating when caribou and ferruginous hawks migrated, our model also identified differences in how the two populations migrated; ferruginous hawks achieved efficient migrations by increasing their movement rates while caribou migration was achieved through significant increases in directional persistence. Our approach is broadly applicable to many animal movement studies. We anticipate that rigorous assessment of migration metrics will aid understanding of both how and why animals move.

## 1 Introduction

Migration is one of the most widespread and important ecological processes within the animal kingdom (Dingle and Drake, 2007; Bauer and Hoye, 2014). The process occurs in countless animal taxa and has evolved convergently many times (Pulido, 2007; Roff and Fairbairn, 2007; Fryxell and Holt, 2013). Owing in part to this convergent evolution, migration is a diverse process, occurring across a wide variety of temporal and spatial scales (Egevang et al., 2010; Hebblewhite and Merrill, 2011; Bohart et al., 2021; Abril-Colón et al., 2022). Understanding how and why animals migrate is important theoretically but understanding where these animals are going and when facilitates effective management (Middleton et al., 2020; Kauffman et al., 2021). As the world undergoes a period of rapid and unprecedented change, the migratory patterns of many animals have changed in response, particularly with respect to their spatial and temporal extent (Hardesty-Moore et al., 2018; Tucker et al., 2018). Recent advances in tracking technology have allowed ecologists to collect animal location data at unprecedented spatial and temporal resolutions, creating opportunities to answer more complex questions pertaining to migration (Kays et al., 2015). This influx of data describes the spatial extents of many animal migrations in detail. The temporal extent of migration is needed for phenological studies but is more difficult to quantify.

Ecologists have designed many approaches to identify the beginning and end of an animal’s migration (Cagnacci et al., 2016; Soriano-Redondo et al., 2020). In some cases, the presence of ecological barriers along an animal’s migratory route make the onset of a migratory period easy to classify without explicit modelling (López-López et al., 2010; Rotics et al., 2018). When these barriers or thresholds are difficult to rigorously define, statistical methods can estimate migration timings. An often-used approach designed by Bunnefeld et al. (2011) relies on net squared displacement (NSD; the animal’s distance from its initial location). The model fits non-linear curves representing different movement strategies (e.g., migratory, nomadic) to explain how an animal’s NSD changes over time. The approach effectively differentiates migratory animals from non-migrants, but it only estimates the “centre” of migration as a parameter, not the beginning or end. Path segmentation analyses focus on dividing a movement path into segments with “change-points” that represent shifts in behaviour (Edelhoff et al., 2016). These models have often been used for identifying area-restricted searching bouts in foraging animals (Weng et al., 2008) but their principles can be extended to identifying migration (Limiñana et al., 2007; Madon and Hingrat, 2014; Mikle et al., 2019; Wolfson et al., 2022). Path segmentation approaches take many forms but broadly, they typically couple a movement metric (e.g., NSD) with a change-point algorithm that identifies changes in the distribution of this metric (Edelhoff et al., 2016). Methods that rely on NSD are sensitive to the animal’s initial location and may break down depending on when data collection began (Singh et al., 2016). First passage time (FPT) is a similar metric that measures the amount of time required for an animal to travel a certain distance, and it has been used to identify changes in movement behaviour on many scales (Johnson et al., 1992; Fauchald and Tveraa, 2003; Le Corre et al., 2014). This distance must be user-defined beforehand, requiring unique assumptions for every dataset (Barraquand and Benhamou, 2008). Complex path segmentation approaches work even when the desired number of segments is not known (Lavielle, 2005; Gurarie et al., 2009; Madon and Hingrat, 2014). Among the plethora of migration models, movement ecologists are still searching for a model that accurately and precisely estimates biologically meaningful parameters that describe when and how animals migrate, with little to no prior knowledge of the system.

Dingle and Drake (2007) provide two separate definitions for migration in individual animals: a persistent period of directionally autocorrelated (or straight) movement, and a period of movement ranging over an exceptionally large spatial extent. Step lengths, the Euclidean distance between two consecutive tracked locations, and turning angles, the angle made by the animal’s turn during three consecutive tracked locations, describe the speed and directionality of a movement track, respectively. Both of these metrics are widely used in movement ecology (Morales et al., 2004; Fortin et al., 2005; van Moorter et al., 2010; Avgar et al., 2016). The first definition of migration suggested by Dingle and Drake (2007) relates to directional persistence, and could be quantified by a change in an animal’s turning angles, while the second definition relates to distance covered and could be quantified by a change in an animal’s step lengths. While many path segmentation models combine these properties into one metric (e.g., NSD or FPT), we suggest that a path segmentation model that identifies simultaneous changes in two metrics (step lengths and turning angles) will allow ecologists to draw more biological context from migration data.

We designed a model that identifies the temporal extent of migration using step lengths and turning angles alone. We hypothesized that migration can be quantified by an abrupt change in an animal’s observed movement speed and directionality for a sustained temporal interval. Unlike most path segmentation approaches, which focus on one all-encompassing movement metric, our model generates distributions for step lengths and turning angles concurrently. We designed a likelihood-based method for identifying the optimal sequence of change-points (e.g., start and end of migration) and used a parametric boot-strapping algorithm to generate confidence intervals for the parameter estimates. Our model works for a diversity of migratory animals sampled at different temporal frequencies, which we display with three case studies: ferruginous hawks (*Buteo regalis*) in the Great Plains of central North America, and barren-ground caribou (*Rangifer tarandus groenlandicus*) and brown bears (*Ursus arctos*) in northern Canada. The inference that can be drawn from this model can have important management implications when applied to additional datasets.

## 2 Methods

### 2.1 The model

Our modelling approach builds on and simplifies existing approaches for estimating the start, end, and intensity of migration. This model only requires information on step lengths and turning angles calculated from a discrete-time sample of an animal’s movement path. If we define **z**_*t*_ = (*x*_*t*_, *y*_*t*_) to be the animal’s recorded location at time *t*, we calculate the step length *r*_*t*_ as follows:

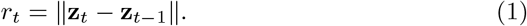

Step lengths are an indicator of the distance an animal travels per time step, and turning angles indicate the directional persistence (or straightness) of movement (Morales et al., 2004). We calculate the turning angle *ϕ*_*t*_ as follows:

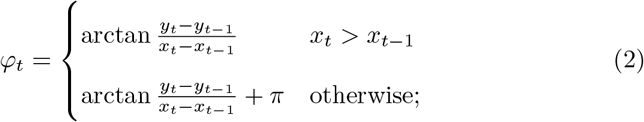

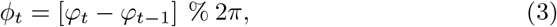

where arctan is the inverse tangent (arc-tangent) function. Applying the modulus operator % ensures that all values are between 0 and 2*π*. Smaller turning angles (closer to 0 or 2*π*) indicate straighter movement.

Step lengths and turning angles are well-studied and can typically be explained effectively using known distributions, which we leverage for our model (Auger-Méthé et al., 2016; Avgar et al., 2016). We hypothesize that an animal’s step lengths follow an exponential distribution at all stages of movement, but during the animal’s migratory stage, the parameter dictating the mean step length increases. We also hypothesize that an animal’s turning angles follow a von Mises distribution, where the angular concentration parameter increases during migration. We assume there exist temporal parameters *t*_1_ and *t*_2_ that signal the start and end of migration, respectively. The likelihood function for any given point **z**_*t*_ incorporates these conditions explicitly with model parameters *t*_1_, *t*_2_, *ρ*_0_, *ρ*_1_, *κ*_0_, and *κ*_1_. During the non-migratory period (*t* < *t*_1_ or *t > t*_2_) the animal’s step length distribution is parameterized by *ρ*_0_ and the animal’s turning angle distribution by *κ*_0_. The parameters *ρ*_1_ and *κ*_1_ represent the additional movement distance and angular concentration incurred during migration, respectively. We define the likelihood function as follows:

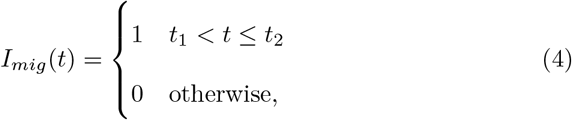

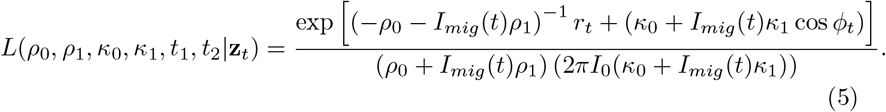

Here, *I*_0_(*κ*) is the modified Bessel function of order 0. The ratio between the animal’s mean step length during and the outside of migration approximates how much more quickly the animal moves when migrating. We denote this quantity 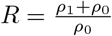.

If necessary, we can also expand the model to account for multiple migratory periods within one dataset. This would necessitate the introduction of additional parameters *t*_3_, *t*_4_, …, *t*_2*c*−1_, *t*_2*c*_ for a model with *c* distinct periods of migratory movement. If *c* > 1, *I*_*mig*_(*t*) would be 1 when *t*_2*n*−1_ < *t* ≤ *t*_2*n*_ for any positive integer *n*. Unique step length and turning angle parameters (*ρ*_2_, …, *ρ*_*c*_ and/or *κ*_2_, …, *κ*_*c*_) for each migratory period could be biologically realistic for some species. For any positive integers *m* and *n*, where *m < n*, the *m*-migration model is nested within the *n*-migration model; this can be verified by setting all *ρ* and *κ* equal to each other and fixing all *t*_*i*_ equal to each other for *i* > *m*.

### 2.2 Parameter estimation

Optimizing the likelihood function (Equation 5) is difficult because the function is not differentiable with respect to temporal parameters *t*_1_ and *t*_2_. The easiest way to solve this problem is to fix all *t*_*i*_ and optimize the model for all *ρ*_*i*_ and *κ*_*i*_. This process can be repeated for every meaningful set of *t*_*i*_ values (there is always a finite number of such combinations with discrete-time data) to find the overall maximum likelihood estimate.

With datasets spanning a wide temporal range (or with *c* > 1), the number of *t*_*i*_ combinations can become problematically large. In these cases, we use an iterative grid-search algorithm to find optimal regions of the likelihood profile quickly, before honing in on those regions with a finer grid. We optimized the *t*_*i*_ over a subsetted grid that only included properly ordered parameter combinations (*t*_*m*_ < *t*_*n*_ if *m* < *n*). The number of grids used and their respective resolution depends on the temporal extent of the data as well as the desired precision with which one hopes to estimate the *t*_*i*_ parameters. The temporal extent of the movement paths varied between datasets but we used a minimum grid size of 1 day for all case studies. Optimizing over a coarse initial grid poses risk of missing global optima but reduces computational times. We started by partitioning the temporal extent of each movement track into 14-day intervals and first found the optimal values of each *t*_*i*_ on this coarser grid. We then identified the *t*_*i*_ combinations that produced the five lowest values of the negative log-likelihood (NLL) function when optimized over *ρ*_*i*_ and *κ*_*i*_; this handles cases when the global optimum may not be near the lowest NLL value along a coarser grid. We then used a finer grid, this time with *t*_*i*_ values spaced 7 days apart, to more thoroughly search these optimal regions. Once again, the 5 lowest NLL values were taken from the 7-day grid for further exploration. We repeated this process with a 3-day grid before finally optimizing along a 1-day grid. By using many grids with a temporal resolution increasing roughly by a factor of two, our algorithm found the optimum much more quickly than using fewer grids, because within each grid there were few *t*_*i*_ combinations to be tested.

We generated a parametric bootstrapping algorithm that estimates 95% confidence intervals for our model’s parameters. We cannot obtain confidence intervals using more standard methods (e.g., Wald-type estimations or likelihood profiles) because the likelihood function includes *I*_*mig*_(*t*), which resembles a step function. The likelihood function is not continuous with respect to the *t*_*i*_ parameters, which shift the position of *I*_*mig*_(*t*). To generate confidence intervals for an individual migration, we simulated random paths with the same size and temporal extent as the true migratory path. The number of random paths necessary to generate consistent confidence intervals varied depending on the dataset. These simulated paths were generated using the likelihood function and parameterized based on the maximum likelihood estimate for each of the model parameters from the true path. We then fit the model to each of these paths independently and used the distribution of the parameter estimates from each random path to obtain confidence intervals (taking the 2.5% and 97.5% quantiles as lower and upper confidence bounds, respectively). The process of re-simulating data according to the estimated parameter values has been used in time-series data for many purposes, including calculating confidence intervals (Dennis and Taper, 1994; Kunst, 2008).

We conducted all data preparation and model fitting using R 4.2.1 (R Core Team, 2021). We obtained maximum likelihood estimates for the *ρ*_*i*_ and *κ*_*i*_ (with the *t*_*i*_ fixed) using the R Template Model Builder (TMB) package (Albertsen et al., 2015; Kristensen et al., 2016).

### 2.3 Simulation analysis

We simulated migratory movement as a series of random step lengths and turning angles, which form a complete path when taken together. Simulation analyses like these allow us to directly compare parameter estimates to “true” parameter values, which cannot actually be identified from animal tracking data. We simulated movement paths over 200 days with 1 observation per day (note that the use of “day” here is for clarity as the time units are arbitrary). Between days 70 and 100, we simulated step lengths from an exponential distribution with a mean step length of 45 km (once again, the spatial units are arbitrary) and turning angles from a von Mises distribution with concentration parameter *κ* = 0.5. Outside of this simulated “migratory period”, these values changed to 5 km and *κ* = 0, respectively. Once we constructed complete movement paths, we randomly removed points such that approximately 150 of the 200 complete “steps” (groups of three consecutive points necessary for calculating turning angles) remained. We accomplished this by removing each point with a probability of 12.5%, which would remove approximately 25% of the complete steps in the data.

We compared our model to three commonly used approaches by fitting them to simulated migratory movement paths. In addition to our model, we fit the NSD regression model from Bunnefeld et al. (2011), the FPT path segmentation model from Le Corre et al. (2014), and a path segmentation approach using daily movement distances (step lengths) from Madon and Hingrat (2014). The two path segmentation approaches use different algorithms for identifying the optimal change-points; Le Corre et al. (2014) use the penalized constant method designed by Lavielle (2005) and Madon and Hingrat (2014) used the Pruned Exact Linear Time (PELT) algorithm designed by Killick et al. (2012). We fit the models to 50 independently simulated migratory paths, all with the same “true” parameters, and calculated the mean bias (estimated *t*_*i*_ - true *t*_*i*_) and mean squared error (MSE; the mean of (bias)^2^ for all 50 samples). The variance of the estimator can be calculated by subtracting MSE from the square of the mean bias, gauging the precision of the model. We used the adehabitatLT R package (Calenge, 2006) to compute FPT time-series and identify changepoints in those time series. We used the changepoint R package (Killick and Eckley, 2014) to run the PELT algorithm. We provide more detail on the implementation of each of these methods in the Appendix.

### 2.4 Case studies

#### 2.4.1 Ferruginous hawks in the Great Plains

Ferruginous hawks are large, migratory raptors found in central Canada and United States (Schmutz and Fyfe, 1987; Schmutz et al., 2008). The shortgrass prairies of southern Alberta, Canada represent the northern edge of this species’s breeding range, and birds breeding this far north make relatively long migrations to the southern Great Plains in the United States (Watson and Keren, 2019). Adult ferruginous hawks were captured at nest sites during the breeding season, using either a dho-gaza net or a bal-chatri trap (Watson, 2020). Captures were limited to nests in which the young had survived at least 10 days. Once captured, the birds were fitted with solar ARGOS/global positioning system (GPS) platform transmitter terminals and solar Groupe Special Mobile (GSM) tags. ARGOS tags recorded a location every 1 hour and GSM tags recorded a location as frequently as every 1 minute (Watson, 2020), so we rarefied each movement track to one location per hour for consistency. Our dataset includes 50 individual hawks tagged on their breeding territories in southeastern Alberta and spans 10 years (2012-2021). The tags also provided estimates of dilution of precision (DOP) in the horizontal and vertical directions for every location. We removed any locations with a DOP over 5 in either the horizontal or vertical directions in preparation for our analysis (Edenius, 1997).

We isolated each individual migration (fall or spring) temporally so we could fit our model with *k* = 1 to them separately. Each hawk was originally tagged on its breeding territory so we used the date at which the first location was received for each individual as the cut-off point between spring and fall. To define a cut-off between the end of fall migration and the beginning of spring migration (i.e., the birds’ arrival at the wintering grounds), we used the date at which the southernmost location was recorded in each year. Once these separations were made, we removed any migrations that were missing a significant section of data, either spatially (any migration containing a location that was further than 400 km away from the previous recorded location) or temporally (any migration containing a 14-day period without any recorded locations). The temporal resolution, or fix rate, of a movement dataset has a significant effect on the results of many movement analyses (Jerde and Visscher, 2005; Thurfjell et al., 2014), so we fit the model to the hawk movement tracks rarefied to 1-hour, 12-hour, and 24-hour fix rates. We bounded *t*_1_ and *t*_2_ such that *t*_2_ − *t*_1_ needed to be greater than 7 days, as anything shorter would represent a biologically unrealistic migration (Watson and Keren, 2019). We also estimated 95% confidence intervals for each individual migration using the parametric bootstrapping method described above. We simulated 100 random paths for each true migratory path. We ran the algorithm multiple times for the same migration and comparing the intervals to ensure that this number of paths produced consistent confidence intervals.

Like many animal species, ferruginous hawks display complex migratory patterns including stopovers and pre-migratory dispersal (Watson et al., 2018; Watson and Keren, 2019). Stopover behaviour is defined as the interruption of migration over some temporal period (Rappole and Warner, 1976) and is very diverse, just like migration itself (Salewski et al., 2007; Evans and Bearhop, 2022; Schmaljohann et al., 2022). Stopovers have many functions and differen-tiating long-term, foraging stopovers from shorter stopovers may be important in identifying critical habitat for migratory species (Green et al., 2002). During fall migration, many ferruginous hawks display long-term stopovers; Watson and Keren (2019) consider these fall movements to be two separate migrations partitioned by the stopover. Ferruginous hawks also frequently embark on premigratory movements, where they disperse from their breeding or winter territory before returning to the same general area (Watson et al., 2018). To evaluate whether our model could statistically identify stopovers and other complexities from the ferruginous hawk data, we compared our model fits with *c* = 1 (one migration) to those with *c* = 2 (two migrations) using Akaike Information Criterion (AIC) and Bayesian Information Criterion (BIC). AIC and BIC compare models by incorporating the maximum log-likelihood estimate along with the complexity of the model, quantified by the number of free parameters. Both metrics are used to select the most parsimonious model but when sample size is large, BIC is more likely than AIC to select models with fewer parameters (Burnham and Anderson, 2004). The model associated with the lowest AIC or BIC value is assumed to be the most parsimonious, and the difference in AIC or BIC between the best model and other models (ΔAIC or ΔBIC, respectively) quantifies how much more parsimonious the best model is. The likelihood ratio test, another common model comparison metric, is not applicable to our model because our likelihood function is not continuous (Self and Liang, 1987).

We performed a short simulation analysis to evaluate whether AIC or BIC could reliably identify stopovers when they were knowingly present (i.e., incorporated into the simulation). We used the same process described in Section 2.3 to generate simulated movement paths but here, our simulation parameters were informed directly by the observed migration timings from ferruginous hawks. See the Appendix for more details on how we parameterized these simulations.

#### 2.4.2 Barren-ground caribou in northern Canada

Caribou are one of the most well-studied species in the animal kingdom (Seip, 1992; Vors and Boyce, 2009; Festa-Bianchet et al., 2011). The many subspecies and ecotypes of caribou exhibit different life history and foraging strategies (Nagy et al., 2011), and the barren-ground caribou herds in the North American Arctic are notable for their migratory behaviour (Lent, 1966; Fleck and Gunn, 1982; Gunn and Miller, 1986; Torney et al., 2018). Our caribou data were collected for the Qamanirjuaq herd, which ranges across Nunavut’s Kivalliq region for much of the spring and summer. This herd moves annually between their more southern winter grounds and their calving and summer ranges further north. Caribou do not always display high inter-annual fidelity to their wintering grounds (Fullman et al., 2021) but, in part due to the gathering of large herds which facilitates social learning, the herd displayed high fidelity to their calving grounds for at least 40 years (Gunn et al., 2012). Pregnant females that arrive on the calving grounds give birth to their calves shortly after, and dramatically reduce their movement for up to two weeks (DeMars et al., 2013; Mallory et al., 2020). Identifying the temporal extent of barren-ground caribou migration has management implications, especially as climate change and anthropogenic modifications to the landscape alter the phenology and availability of their food resources (Chen et al., 2018; Mallory et al., 2020). Many efforts have been made to identify these timings in other herds (DeMars et al., 2013; Le Corre et al., 2014; Torney et al., 2018).

We fit the migration model with *c* = 1 to data describing the spring migrations of barren-ground caribou. Caribou were pursued via helicopter and immobilized via net-gunning, before being fitted with a GPS collar (Mallory et al., 2020). Following approved protocols, caribou were collared between 2006 and 2016 and in total, we included 35 adult females in the dataset, of which 22 were tracked for more than 1 year. We isolated each individual year and sub-setted the data such that any locations after July 1 of that year were omitted. We chose this date because it is after the calving period (Mallory et al., 2020) but earlier than the onset of fall migration (Le Corre et al., 2017). The fix rates of each individual in the dataset varied from 1 hour to 1 day, so we rarefied all the data to a 1-day fix rate for consistency. Similarly to the ferruginous hawk dataset, we removed any migrations with significant spatial (150 km between two consecutive recorded locations) or temporal (any 14-day period without recorded locations) gaps from our dataset.

Caribou migrations are well-studied, so we compared migration timings from our model to those identified by an existing, commonly used approach (Bunnefeld et al., 2011). This approach involves fitting a non-linear (specifically, logistic) curve to an animal’s net squared displacement as a function of time. When only modelling one migration (and not the return back to the wintering grounds), the NSD model contains three parameters (see Equation 3 of Bunnefeld et al., 2011): the asymptotic NSD value *δ*, the peak or “centre” of migratory movement *θ*, and the quarter-duration of migration *ϕ*. The beginning and end of migration are estimated based on the estimates of *θ* and *ϕ*:

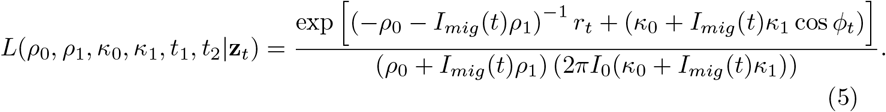

To leverage the wealth of information on barren-ground caribou migration and calving, we fit a second version of our model that incorporated *a priori* knowledge of calving phenology. Caribou calving can be identified from movement data because when females give birth to calves, they greatly reduce their movement rate (DeMars et al., 2013). We used the broken-stick regression method developed by DeMars et al. (2013) to identify the beginning of the calving period, which also conveniently signaled the end of spring migration. With this information to our advantage, we fit the migration model again, but this time we fixed *t*_2_ to be equal to the date at which calving began, and only optimized along *t*_1_. This reduces the complexity of our grid and allowed us to optimize *t*_1_ immediately along the 1-day timescale, without need for the iterative grid-search method described in Section 2.2. We visually compared the results from the NSD model, our model with no *a priori* information, and our model with *a priori* information to determine which model was most effective in identifying meaningful shifts in caribou movement behaviour.

#### 2.4.6 Brown bears in northern Canada

Brown bears are opportunistic omnivores with a wide distribution across North America, Europe, and Asia (Pasitschniak-Arts, 1993). Brown bears in the Canadian Arctic are unique in comparison to their conspecifics worldwide, exhibiting many adaptations to harsh environmental conditions (Edwards and Derocher, 2015). Brown bears are not considered migratory, but bears living in the Mackenzie River Delta region of northern Canada display annual home range shifts (Edwards et al., 2009) and perform temporally oriented navigations to food resources visited a year prior (Thompson et al., 2022). We used brown bear movement data from the Mackenzie Delta to evaluate if our model would identify any patterns in what biologists view as a non-migratory species. Brown bears were captured, immobilized, and equipped with GPS collars between 2003 and 2006 (Edwards et al., 2009). These collars were set to record GPS locations at a 4-hour fix rate. Brown bears in the Canadian Arctic spend up to 6-7 months of the year in a den where they hibernate (Halloran and Pearson, 1972; Nagy et al., 1983; McLoughlin et al., 2002). In total, we included 30 bears (24 females, 6 males) in our analysis.

Given the broad definitions of migration (Dingle and Drake, 2007) and the simplicity of our model, we saw value in searching for population-level trends in periods of high-intensity movement within the brown bear dataset. We fit the model with two migratory periods (*c* = 2) to every individual year in the dataset (many individuals had more than one complete year of data), under the assumption that bears would need to exhibit at least two periods of high-intensity movement to complete their theoretical migratory cycle. We then collectively analyzed the population-level distribution of *t*_*i*_ for each model to determine whether any trends persisted.

## 3 Results

All parameter estimates and confidence intervals (from simulated and real migrations) can be found in Supplementary File 1, which is available at github.com/pthompson234/migrationmodelling.

### 3.1 Simulation analysis

Our model was more precise and accurate than other migration modelling approaches when fit to simulated data (Table 1). When the number of changepoints (here, two) is known, our model estimated the temporal duration of migration to within 1 day, on average. Applying the PELT algorithm (Killick et al., 2012; Madon and Hingrat, 2014) to daily distance time-series calculated from the simulated paths produced similarly accurate and precise estimates of migration timing. The NSD model does not explicitly estimate the beginning and end of migration as parameters but we calculated these quantities based on the parameters the model does estimate (Equation 7). These estimates displayed low bias and high variance, suggesting high accuracy and low precision. Estimating change-points with the penalized constant method applied to first passage time data produced high bias and MSE.

**Table 1.** Bias (parameter estimate - true value) and mean squared error (MSE; bias squared) values for migration timing parameters *t*_1_ and *t*_2_, estimated by fitting four different models to 50 randomly simulated migratory movement paths. Variance is calculated as the difference between MSE and (bias)^2^. We compared our model to the net squared displacement (NSD) approach developed by Bunnefeld et al. (2011), the first passage time (FPT) approach from Le Corre et al. (2014), and the PELT algorithm used by Madon and Hingrat (2014).

| Model | $t_1$ Bias | $t_1$ MSE | $t_1$ Variance | $t_2$ Bias | $t_2$ MSE | $t_2$ Variance |
| --- | --- | --- | --- | --- | --- | --- |
| Our model | -0.1 | 2.7 | 2.7 | 0.5 | 1.4 | 1.2 |
| NSD | 4.1 | 182.4 | 165.6 | -7.9 | 511.1 | 448.7 |
| FPT | -10.4 | 435.7 | 327.5 | 22.0 | 1231.6 | 747.6 |
| PELT | 0.6 | 4.6 | 4.2 | -3.2 | 45.6 | 35.36 |

### 3.2 Ferruginous hawks: stopovers and fix rates

We identified 99 unique ferruginous hawk migrations (35 fall, 64 spring). Our model precisely identified the beginning and end of these migratory movements (a specific migration is shown in Figure 1). Ferruginous hawks rapidly increased their step lengths during migration but did not display as much change in their directionality. The average value of *R*, which approximates the proportional increase in animal step lengths during migration, was approximately 11.02 for ferruginous hawks sampled at a 1-hour fix rate. Before and after migration, ferruginous hawk step lengths averaged 1.34 km (the mean of all *ρ*_0_ estimates for each migration), and this increased by 10.62 km (the mean *ρ*_1_ estimate) during migration. Estimates of *κ*_0_, which quantified movement directionality outside of migration, were frequently 0, which would suggest a turning angle distribution that was either very close to uniform or not centred at 0 (Supplementary File 1). The median 95% confidence interval width for all six of our model parameters (0.39 days, 0.33 days, 0.10 km, 2.99 km, 0.06, and 0.18 for *t*_1_, *t*_2_, *ρ*_0_, *ρ*_1_, *κ*_0_, and *κ*_1_, respectively) suggests that all model parameters are estimable (Supplementary File 1). Independent runs of the parametric bootstrapping algorithm produced similar results for the same data. The largest confidence interval width for either timing parameter (*t*_1_ or *t*_2_) for any individual was 26.88 days, and no other confidence interval for these parameters was wider than 8 days.

**Figure 1:**
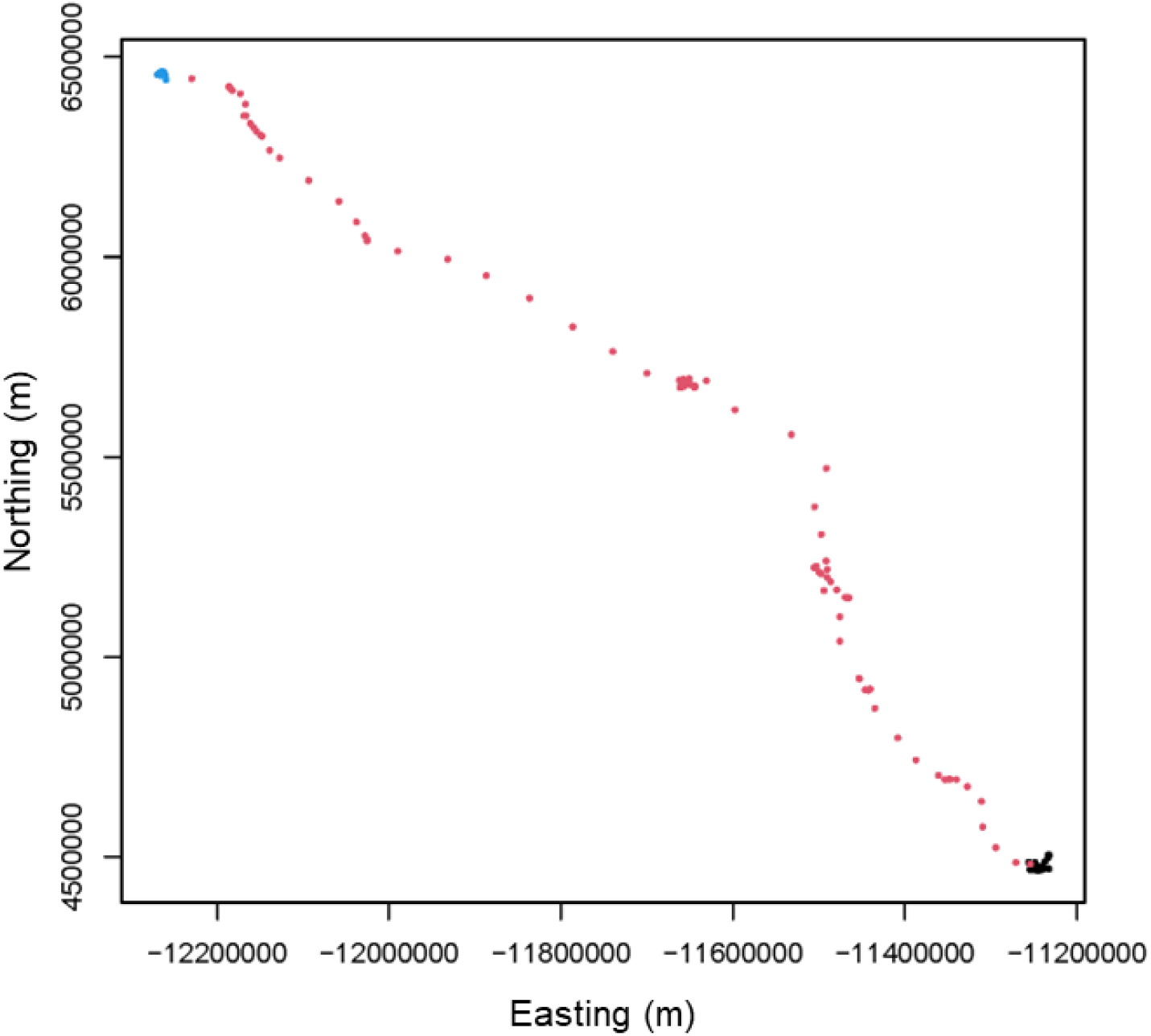
Movement path of a ferruginous hawk (hawk ID 192a; spring 2014) performing a spring migration from its wintering grounds in the southern United States to its breeding territory in southeastern Alberta, Canada. Black dots represent the wintering grounds, blue dots represent the breeding grounds, and red dots represent the migratory period as fit by our model.

The *c* = 2 model identified the timing and location of stopovers and pre-migratory movements in ferruginous hawks. Fall migrants frequently exhibited stopover behaviour, sometimes migrating for *>*500 km before drastically and temporarily reducing their movement rates. The *c* = 1 model occasionally identified only one portion of the fall migration in these cases, but sometimes ignored the stopover altogether (Figure 2). The *c* = 1 model typically ignored pre-migratory movements but sometimes included them as part of the migration (Figure 3). The *c* = 1 model was identified as less parsimonious than the *c* = 2 model when compared with AIC and BIC when these behaviours were present (Supplementary File 1). For example, the migration depicted in Figure 2 had much lower AIC and BIC values with the *c* = 2 model (ΔAIC = 1879.0; ΔBIC = 1866.6, with 1-hour fix rate). The results were similar for the migration depicted in Figure 3 (ΔAIC = 104.6; ΔBIC = 93.6, with 1-hour fix rate). Our simulation analysis provides further support for AIC and BIC as consistent identifiers of stopovers (Supplementary File 1).

**Figure 2:**
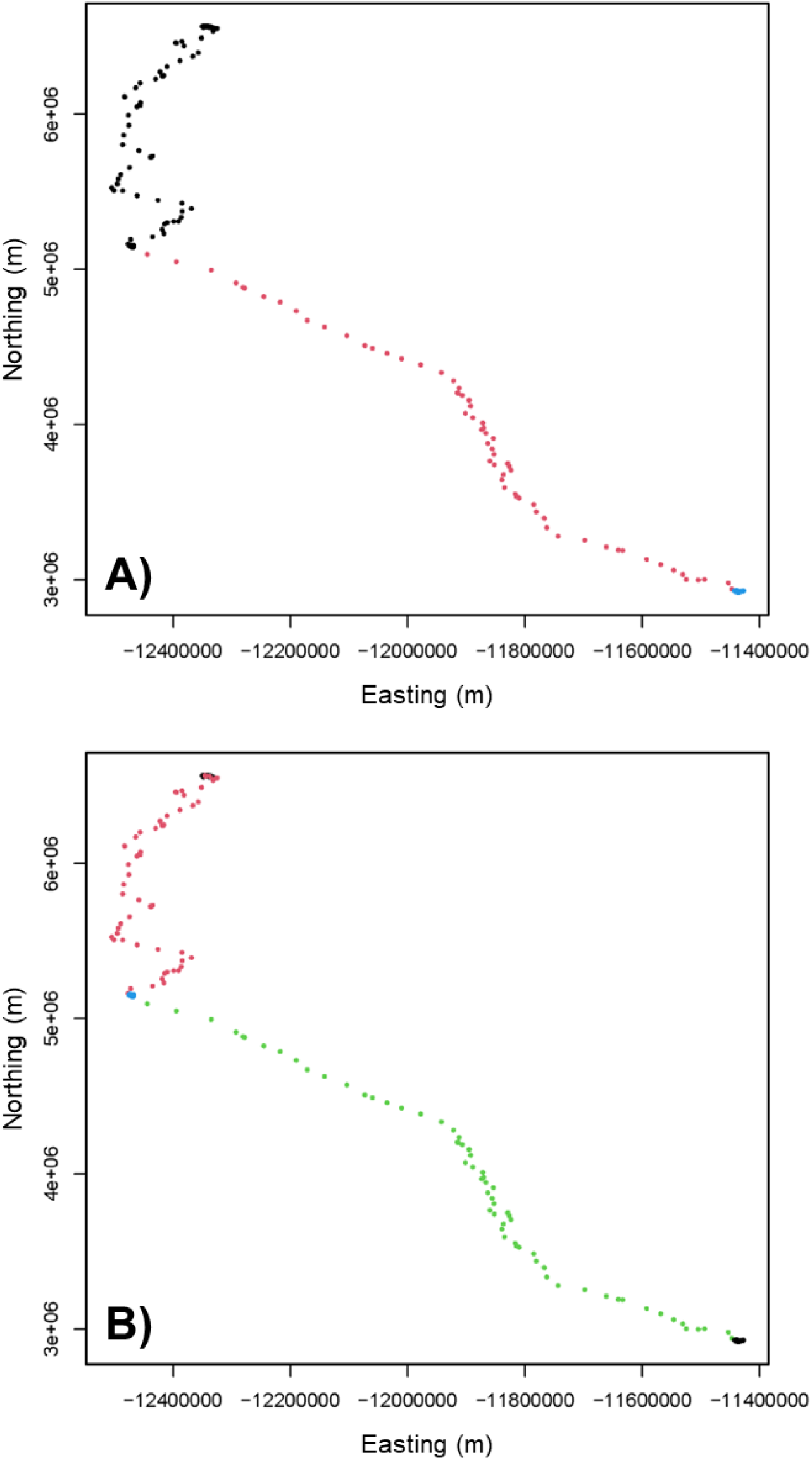
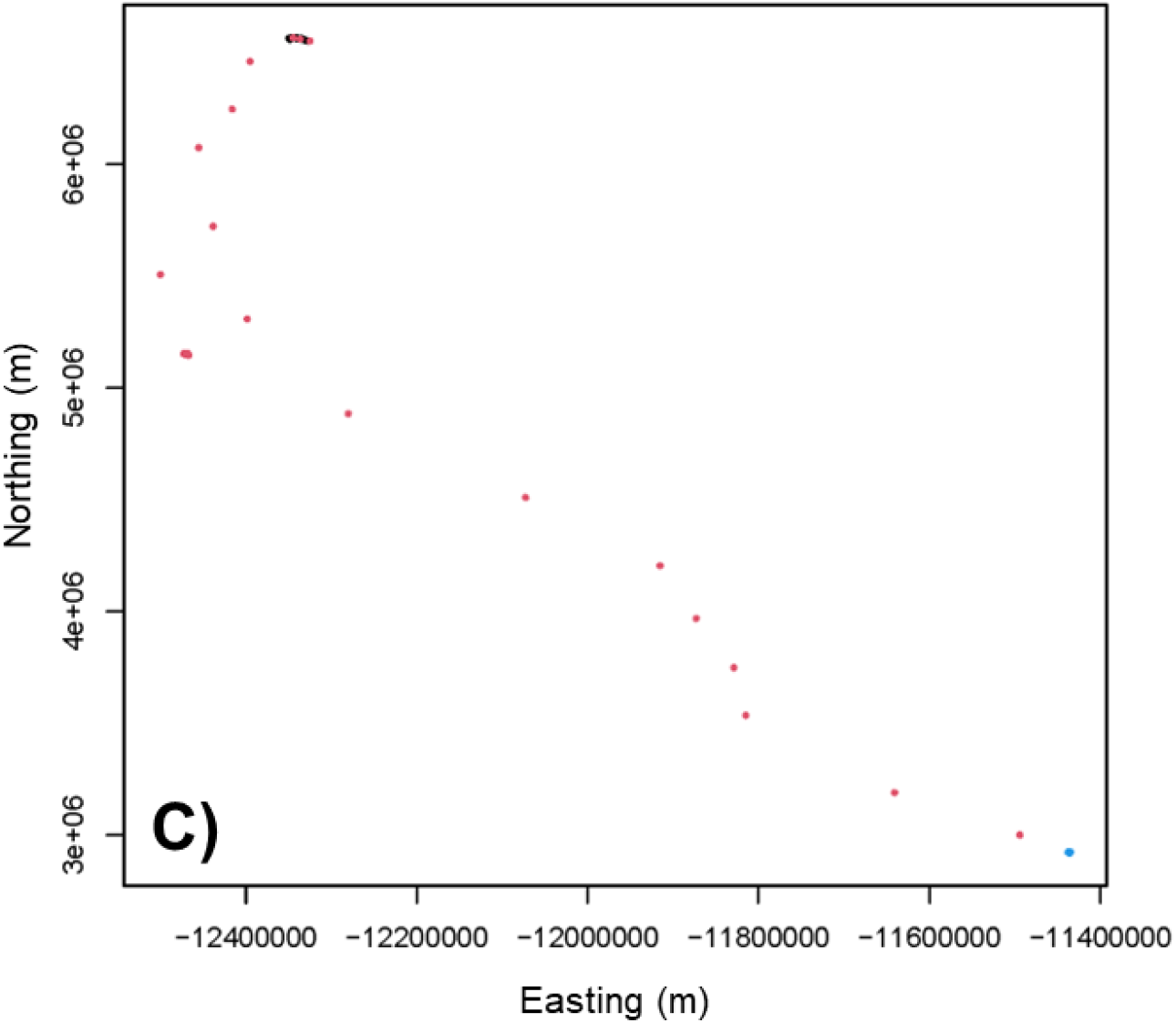
Movement path of a ferruginous hawk (hawk ID 196a; fall 2013) performing a fall migration from its breeding territory in Alberta, Canada, to its wintering grounds in the United States, including a stopover. Panel A) represents the model fit from the *k* = 1 model to the 1-hour fix rate data. Here, black dots represent the breeding grounds, blue dots represent what the model identifies as the wintering grounds, and red dots represent the migratory period. Panel B) represents the model fit from the *k* = 2 model to the 1-hour fix rate data. Here, black dots represent the breeding and wintering grounds, blue dots represent the stopover site, and red and green dots represent the first and second migrations, respectively. Panel C) represents the model fit from the *k* = 1 model to the 24-hour fix rate data. Each point is coloured similarly to in Panel A).

**Figure 3:**
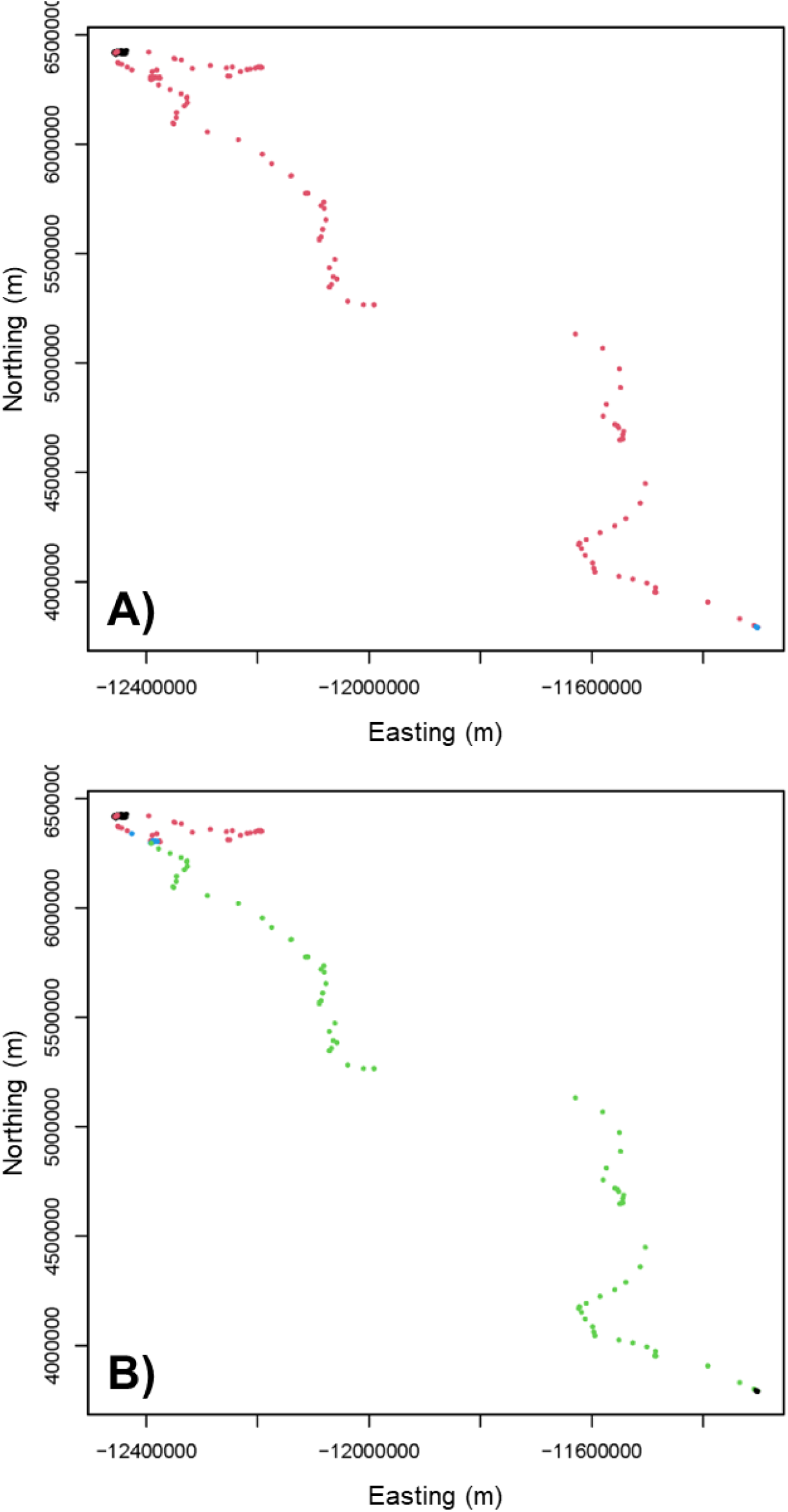
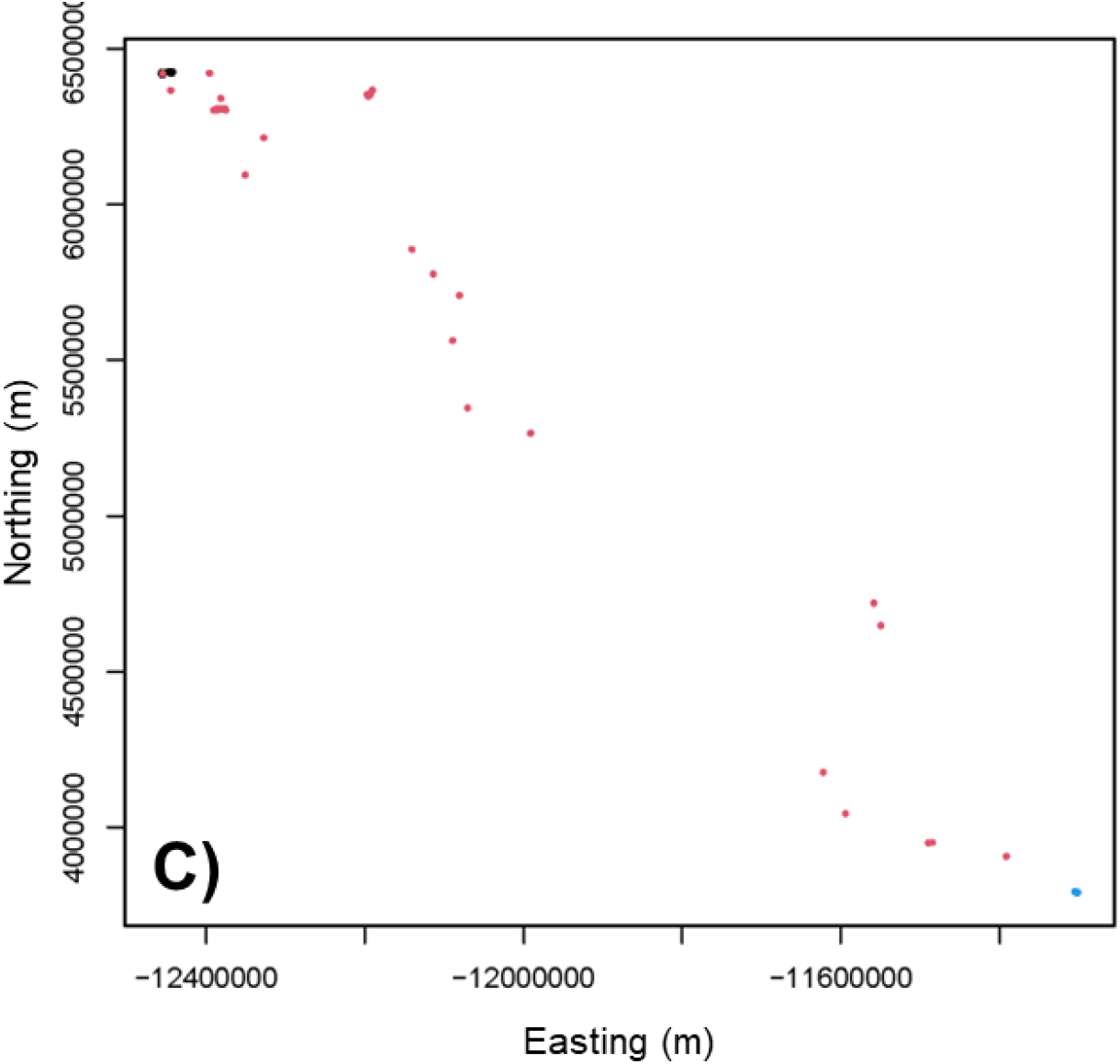
Movement path of a ferruginous hawk (hawk ID 191a; fall 2016) performing a fall migration from its breeding territory in Alberta, Canada, to its wintering grounds in the southern United States, including a pre-migratory movement. Canada. Panel A) represents the model fit from the *c* = 1 model to the 1-hour fix rate data. Here, black dots represent the breeding grounds, blue dots represent what the model identifies as the wintering grounds, and red dots represent the migratory period. Panel B) represents the model fit from the *c* = 2 model to the 1-hour fix rate data. Here, black dots represent the breeding and wintering grounds, blue dots represent the period between pre-migration and migration, and red and green dots represent what the model identifies as the first and second migrations, respectively. Panel C) represents the model fit from the *c* = 1 model to the 24-hour fix rate data. Each point is coloured similarly to in Panel A).

Varying the fix rate of our data did not significantly affect the estimation of ferruginous hawk migration timings but did affect the estimates for step length and turning angle parameters. In some of the migrations with long-term stopovers or pre-migratory movements, the *c* = 1 model estimated different *t*_*i*_ values at different fix rates (e.g., Figure 2). For *c* = 2, temporal parameter estimates were more consistent (Supplementary File 1). Estimates for *ρ*_0_ and *ρ*_1_ were unsurprisingly highest at long fix rates. The mean values of *ρ*_0_ and *ρ*_1_ were 5.18 km and 131.40 km, respectively, when we fit the model to the 24-hour data. The hawks’ proportional increase in movement speed, *R*, appear larger at higher fix rates (59.4 with the 24-hour data and 39.8 for the 12-hour data). Estimates for *κ*_0_ were very close to, if not exactly, 0 at all fix rates. The mean estimate for *κ*_1_ increased from 0.48 with 1-hour fix rates to 1.34 with 24-hour fix rates.

### 3.3 Caribou: incorporating calving phenology

After filtering the caribou data, we retained 57 individual spring migrations to which we fit the *c* = 1 migration model. In many cases, the model identified a biologically reasonable migratory period. The NSD model did the same but often failed to precisely estimate the beginning and/or end of migration. In the example from Figure 4, the NSD model estimated a migration that started 7 days later and ended 2 days earlier than our model. From visual inspection, it appears that our model correctly captures more of the linear migratory component than the NSD model. However, for some caribou-years, our model misidentified a period of sustained movement on the wintering grounds as migration, rather than identifying the spring movement to the calving grounds (Figure 5, Supplementary File 1). In many of these cases, the NSD model picked a more appropriate midpoint but still failed to properly characterize the beginning and end of migration (Figure 5). By estimating *t*_2_ from broken-stick regression models fit to caribou step length data (DeMars et al., 2013), the model consistently identified the biologically relevant spring migratory period. Supplying the model with this additional information remedied the problematic model fits like the example displayed in Figure 5, and allowed us to pinpoint the day at which each caribou began migrating.

**Figure 4:**
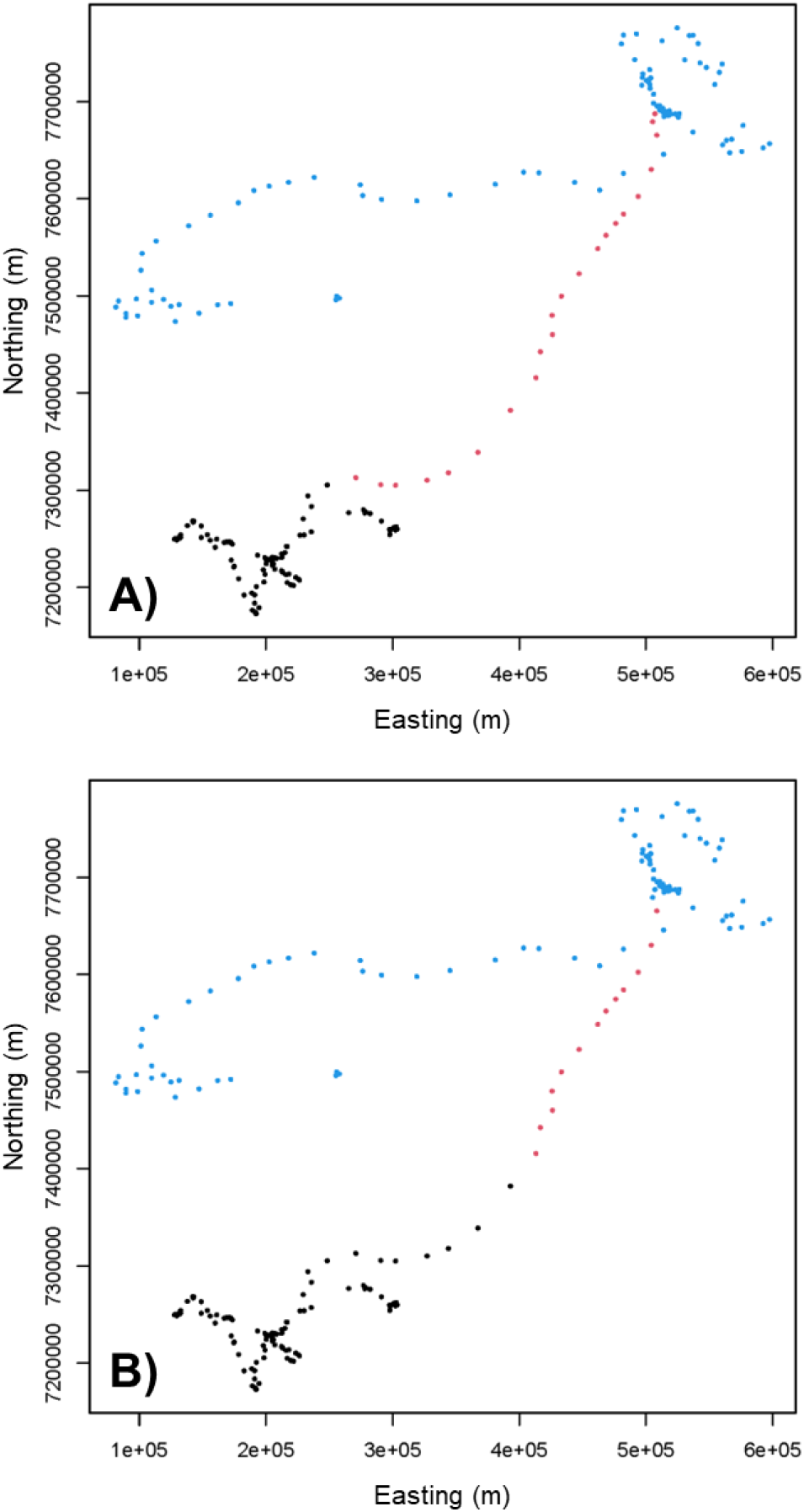
Movement path for a female caribou (caribou ID BL0560413; year 2016) in the Qamanirjuaq herd in Canada. Here, panel A) illustrates the migration timing estimates from our model, and panel B) illustrates the migration timing based on the NSD method developed by Bunnefeld et al. (2011). In both panels, black dots represent the wintering grounds, red dots represent migratory movement, and blue dots represent post-migratory movement. The calving grounds can be visually identified as a tightly packed clump of points just east of the end of migration.

**Figure 5:**
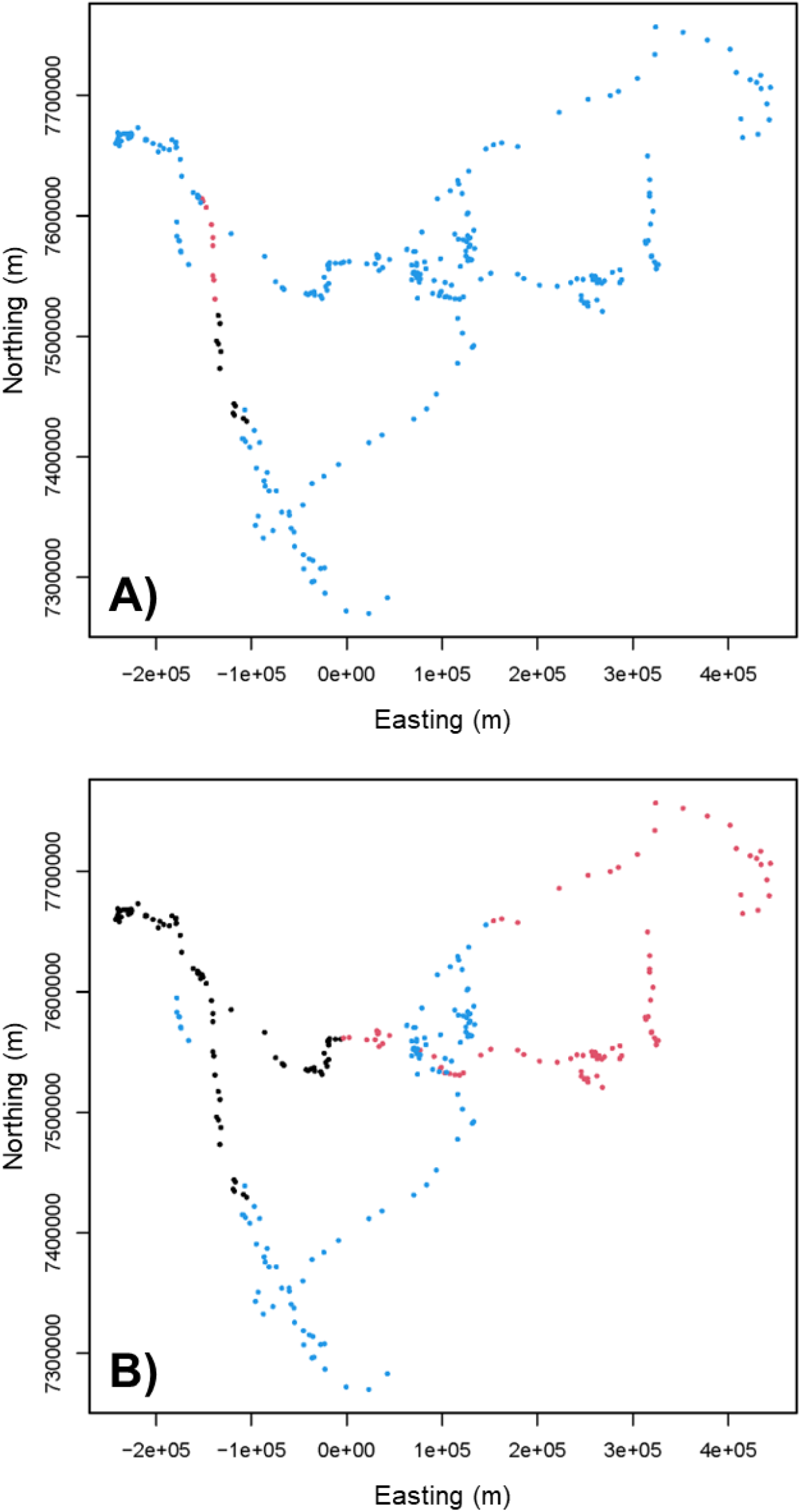
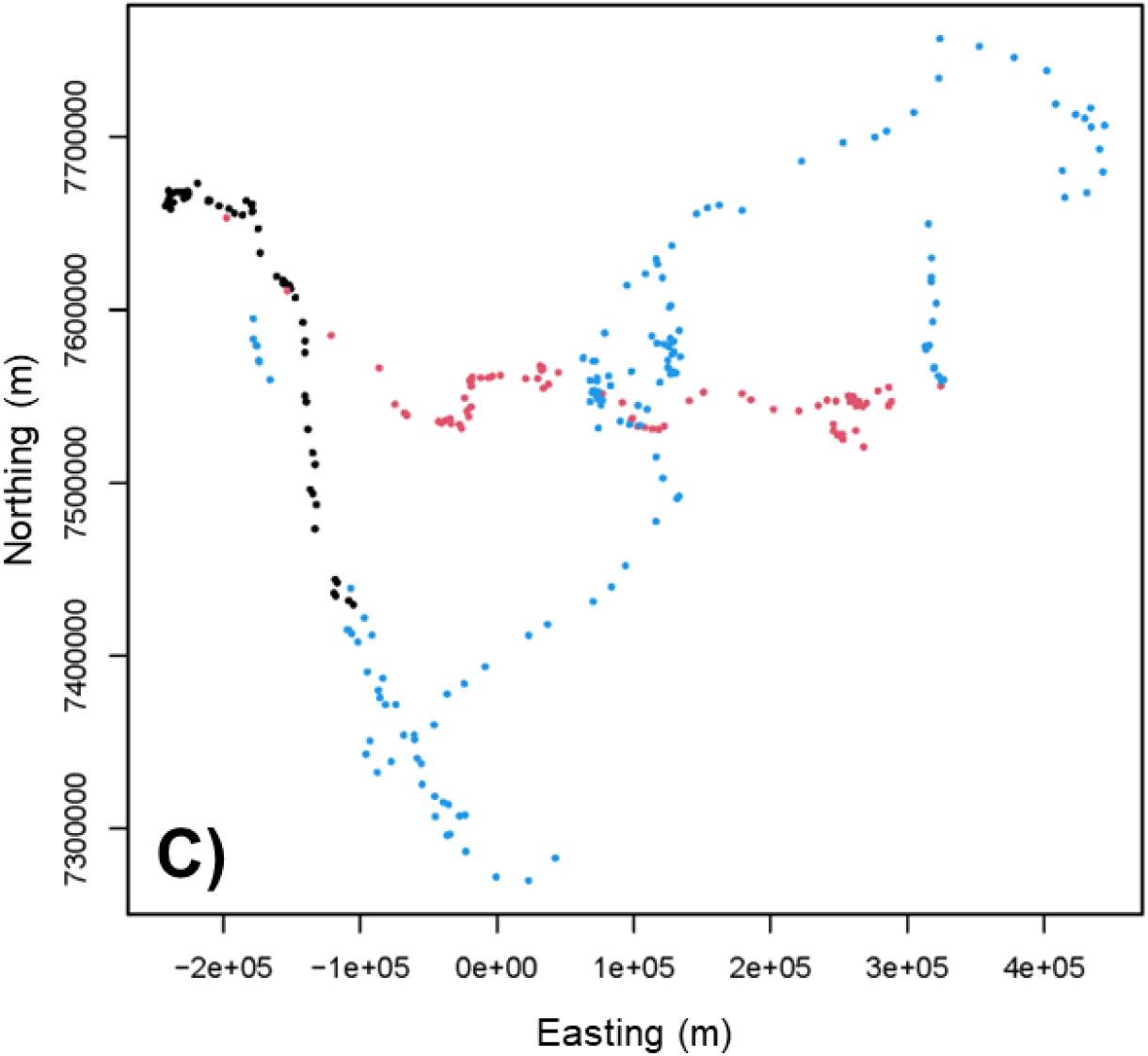
Movement path for a female caribou (caribou ID QM0830508; year 2011) in the Qamanirjuaq herd in Canada. In all three panels, black dots represent the wintering grounds, red dots represent migratory movement, and blue dots represent post-migratory movement. The calving grounds can be visually identified as a tightly packed clump of points just east of the end of migration. Here, panel A) illustrates the migration timing estimates from our model without *a priori* knowledge of calving phenology, and panel B) illustrates the migration timing based on the NSD method developed by Bunnefeld et al. (2011). Panel C) displays the fit from our model after using the technique developed by DeMars et al. (2013) to estimate *t*_2_, the onset of calving and end of spring migration.

Caribou did not increase their speed as much as ferruginous hawks during migration, as the mean value of *R* was 3.95 (Supplementary File 1). However, 49 of the migrations displayed significantly higher directional persistence on migration, with 95% confidence intervals for *κ*_1_ excluding 0. The median confidence interval width for *t*_1_, *ρ*_0_, *ρ*_1_, *κ*_0_, and *κ*_1_ were 16.5 days, 1.21 km, 9.43 km, 0.37, and 1.87, respectively. Individuals with low estimated values of *ρ*_1_ and *κ*_1_ were an exception, because paths simulated by our bootstrapping technique exhibited little change during the migratory period (Supplementary File 1). The six migrations with a 95% confidence interval for *t*_1_ that was wider than 100 days all satisfied *κ*_1_ *<* 0.6 (compared to the mean value of 2.05), and five of the six satisfied *ρ*_1_ *<* 4 km (compared to the mean value of 9.14 km).

### 3.4 Brown bears: application to non-migrants

We fit the *c* = 2 model to 42 different bear-years and could not identify any trends throughout the population. According to the model results, 29 of the individuals spent over half of their active season “migrating”, and 11 “migrated” for over 75% of the active season (Supplementary File 1). In other individuals, the duration of one or both of the theoretical migratory periods was 7 days or shorter. While the model appears to have identified periods in which brown bears moved more quickly and/or less tortuously for a number of days or weeks, there was no consistency within the population as to when these periods took place or how long they lasted (Figure 6).

**Figure 6:**
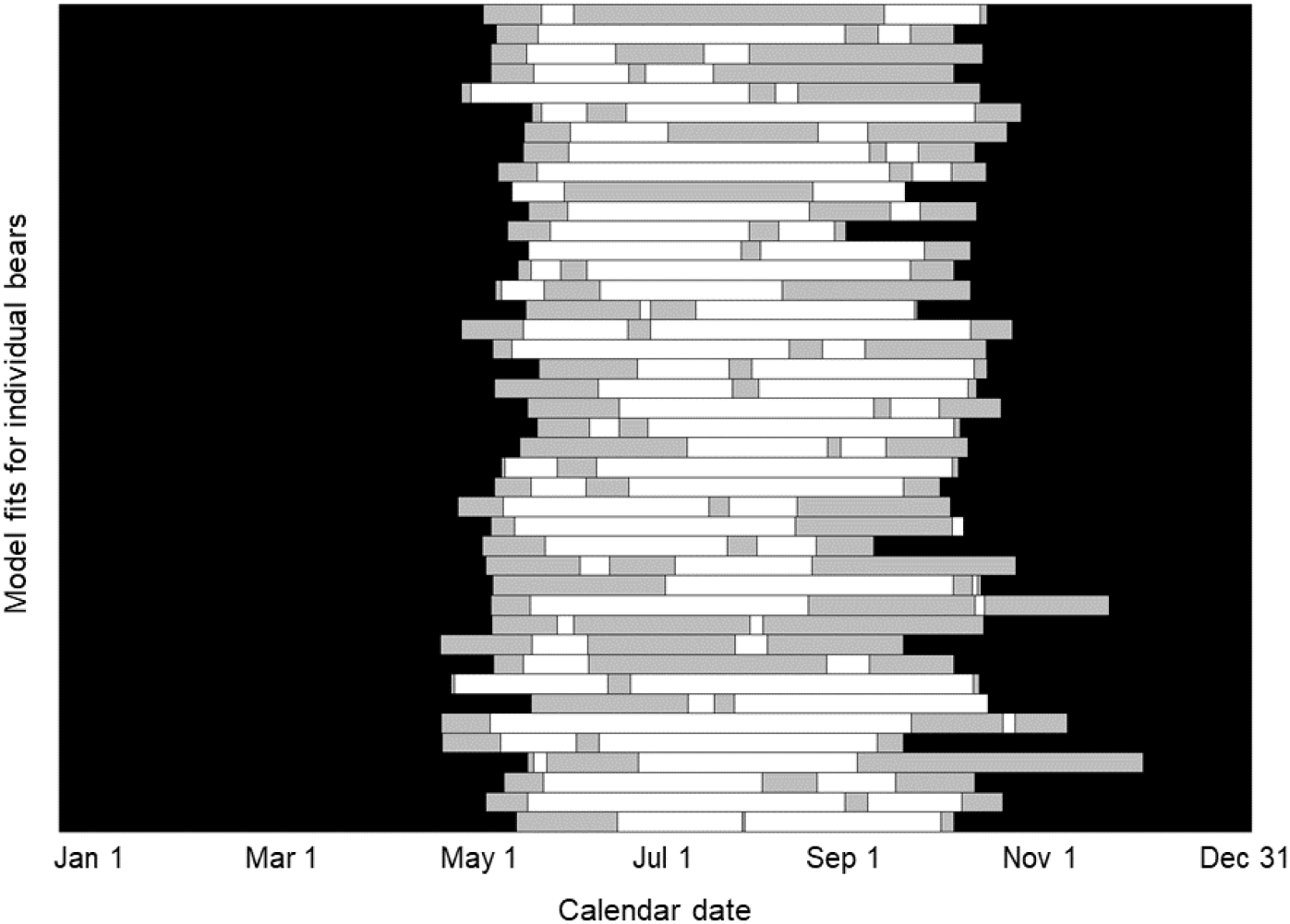
Distribution of theoretical Mackenzie Delta brown bear migratory patterns for 30 individuals (46 bear-years). Portions of the graph shaded in black represent the denning period each year, gray-shaded portions represent non-migratory portions (either before, between, or after both the migrations), and white portions represent theoretical migrations identified by the *c* = 2 migration model.

## 4 Discussion

Identifying when animals migrate is crucial to understanding and predicting changes in migration phenology as a response to climate change (Hardesty-Moore et al., 2018). We designed a model that synthesizes these existing approaches to estimate the timing of an animal’s migration. Our model out-performed competing approaches when fit to simulated data (Table 1), and also identified biologically reasonable timings for migratory mammals and birds (Figures 1 and 4). We failed to identify any significant trends in migration-like behaviour for animals that are not considered migratory (Figure 6). The model relies on step lengths and turning angles, which are ubiquitous in animal movement modelling and can be calculated easily (Morales et al., 2004). As a result, the parameters we estimated with the model have direct biological inter-pretations that help describe multiple facets of migration. Our model explicitly estimated the beginning and end of migratory movement and was more accurate than commonly used methods, which we demonstrated using simulated migratory paths for which “true” parameter values were known. When fit to animal tracking data, our model estimated biologically reasonable (e.g., Figure 1) timings with high certainty, according to our 95% confidence intervals. The model does not require *a priori* biological knowledge of a system to identify timings, but is also flexible to include this information if it improves results (Figure 5). Applying our model to other migratory systems will further identify how it can most effectively be used.

In addition to the beginning (*t*_1_) and end (*t*_2_) of migration, our model estimated parameters that quantify exactly how an animal’s movement changes during migration. By combining step lengths and turning angles to identify migration in ferruginous hawks and barren-ground caribou, our model facilitated a connection between parameter estimates and the biological definitions of migration for these species. We used 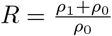 and *κ*_1_ − *κ*_0_ to quantify increases in speed and directionality during migration, respectively, as they can be easily compared between species. Ferruginous hawks moved in a more directed manner during migration, but also moved much more quickly (Supplementary File 1). Migratory ferruginous hawks typically moved 10 times as far during migration than they would otherwise, as the second definition of migration provided by Dingle and Drake (2007) postulates. Migratory barren-ground caribou dramatically increased the directional persistence of their movement during migration and did not “speed up” as much as the ferruginous hawks did (Supplementary File 1). These migrations resembled the “undistracted and persistent” definition of migration from Dingle and Drake (2007).

Fix rate did not dramatically affect *t*_*i*_ estimates for the ferruginous hawk dataset, but estimates of the *ρ*_*i*_ and *κ*_*i*_ varied. It is unsurprising that step lengths become longer as fix rates become longer, but notably, *ρ*_*i*_ estimates did not scale linearly with fix rate. In other words, the estimated value of *ρ*_0_ with data sampled at 24-hour fix rates was less than 24 multiplied by the corresponding *ρ*_0_ at 1-hour fix rates. We advocate for using *R* to compare how different animals migrate but this quantity also varies with fix rate, so this must be controlled before comparing different datasets (e.g., by subsampling, as we did here). Sampling data to coarser fix rates omits the tortuosity of movement at smaller scales, so longer step lengths underestimate movement speed (Postlethwaite and Dennis, 2013). Longer fix rates also produce straighter turning angles (Jerde and Visscher, 2005), which our model quantified with larger *κ*_*i*_ values.

We applied parametric bootstrapping to our model to obtain confidence intervals, and in most cases these intervals suggest high certainty in our parameter estimates (Supplementary File 1). If *ρ*_1_ and *κ*_1_ are both small, the simulated path would not change much during the simulated “migratory” period, making the change-point difficult to estimate. Typically, this is not the case in migratory animals, although some of the caribou paths displayed this result. When this was the case, 95% confidence intervals for all parameters were concerningly wide (Supplementary File 1). These migrations were also difficult to estimate with the NSD model, which typically produced biologically suspicious results in these cases. Accounting for uncertainty in path segmentation models is difficult because of the focus on dividing movement into discernible sections (Edelhoff et al., 2016). We hope our application of parametric bootstrapping to this problem encourages ecologists to incorporate the uncertainty of change-point algorithms into their analyses.

Unlike simple threshold-based approaches, our model does not require any biological knowledge of the tracked animals. Nevertheless, incorporating *a priori* information about an animal’s movement ecology is easy because of our grid-based temporal optimization approach. We directly controlled the set of *t*_*i*_ values included in our parameter space, allowing for the removal of biologically unreasonable timings. This process has the added benefit of reducing computational time. By estimating the beginning of the calving period in adult female caribou (DeMars et al., 2013), we removed an entire parameter (*t*_2_) from optimization. The discrete-time nature of the data for which our model is intended along with our optimization algorithm allows for effective manipulation and improvement of the analysis, but only if necessary.

AIC and BIC were both effective at separating ferruginous hawk migrations with stopovers or pre-migratory periods from those without. Here, AIC and BIC produced similar results but this may not hold at different sample sizes, and picking between AIC and BIC can be a complex problem (Burnham and Anderson, 2004). We note that our model was not effective at distinguishing between stopovers and pre-migratory periods, and this may require additional biological input. We did not test a version of the model with *c >* 2. The computational time allotted by the grid-search optimization algorithm increases exponentially with *c*. We are thus unsure if AIC and BIC are reliable when *c* is larger. Other path segmentation methods (including the PELT algorithm) use more complex techniques for identifying the optimal number of segments that make them extremely useful when the number of segments is unknown (Lavielle, 2005; Gurarie et al., 2009; Killick et al., 2012). In migratory animals, when the number of segments typically is known, our model outperformed other path segmentation approaches, but these competing models are likely more suitable otherwise.

Our model is intended to be fit to movement data for one individual, but fitting the model to several individuals in the same population characterizes the variation within that population. While analyzing the distribution of parameter values across a set of individuals (or individual migrations) is fairly straightforward (Figure 6), there is also an opportunity to regress our parameter estimates (particularly, the *t*_*i*_ parameters) against covariates. Animal populations display high individual variation with respect to their migratory behaviour (Hanski et al., 2004; Jesmer et al., 2018; Merkle et al., 2019; Byholm et al., 2022), and inter-annual variation in an animal’s environment can cause an individual’s migratory paths to vary from year to year (Tucker et al., 2018; Mallory et al., 2020; Franklin et al., 2022). Similarly, the effects of habitat modification and/or associated disturbance factors could also be assessed. With the appropriate environmental data, our model could be used to explore these patterns for many animal populations.

Our model achieved the sought-after goal of determining when animals begin and end their migrations. By parameterizing time-dependent step length and turning angle distributions, we generated results that are easy to interpret biologically. Migration incurs an elevated risk to the negative effects of anthropogenic global change. Specifically, many animals are arriving at their breeding grounds earlier to capitalize on global warming-induced advances in green-up and prey availability (Haest et al., 2018; Mallory et al., 2020). Many ecologists expect (or are already observing) changes in when, where, and how animals migrate (Wilcove and Wikelski, 2008; Tucker et al., 2018). Our model provides unbiased, quantitative information on all three of these characteristics.

## Appendix

**Simulation analyses**

### Model comparison analysis

We simulated migratory movement paths and fit four models to each path. For each path we calculated net squared displacement (NSD) at each time *t* as the Euclidean distance between the animal’s location at time *t* and the animal’s initial location. We fit the model from Bunnefeld et al. (2011) to the NSD time-series. Bunnefeld et al. (2011) model NSD as a function of time using a non-linear regression with three paramters: the intensity of migration *θ*, the temporal “centre” of migration *θ*, and the “quarter-duration” *ψ*. The model takes a logistic form:

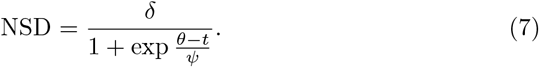

We fit the model using the “nls” function from the R software. As the name of the parameter suggests, we can calculate the beginning and end of migration using *θ* − 2*ψ* and *θ* + 2*ψ*, respectively (Bunnefeld et al., 2011).

We calculated first passage time (FPT) to fit the model from Le Corre et al. (2014), which identifies change-points with the penalized constant algorithm (Lavielle, 2005). We calculated FPT at any time *t* as the amount of time required for the animal’s NSD to exceed some radial threshold *D*. The radius needs to be decided *a priori*, but we can identify the optimal *D* by maximizing the variance of log FPT for a sequence of candidate radii (Fauchald and Tveraa, 2003). We selected our optimal *D* for each simulation separately, choosing among multiples of 5 km increasing to 100 km (5, 10, …, 95, 100). Once we picked *D* and calculated the FPT time series for each simulation, we applied the penalized constant method to identify the optimal FPT change-points (Lavielle, 2005). We used the “lavielle” R function from the adehabitatLT package to run the penalized constant algorithm, following the methods of Le Corre et al. (2014) with the exception of fixing the number of change-points.

We applied the Pruned Exact Linear Time (PELT) algorithm to time-series data representing daily distance travelled for each simulation (for our data, these quantities were equivalent to step lengths). We followed the methods of Madon and Hingrat (2014) as closely as possible, applying the “cpt.var” function from the changepoint R package to the daily distance time-series for each simulation. Once again, the difference between our analysis and the analysis from Madon and Hingrat (2014) was that we fixed the number of change-points at two.

### Simulating hawk-like migrations

We simulated movement paths intended to resemble ferruginous hawk migrations with and without stopovers. We then fit the *c* = 1 (one-migration) and *c* = 2 (two-migration) versions of the model to all these simulated paths and compared the parsimony of each model type using Akaike Information Criterion (AIC) and Bayesian Information Criterion (BIC). We simulated our paths as a series of 1-hour movements over a 200-day period, in line with the hawk data. This gave us 4800 step lengths and turning angles from which we iteratively constructed a movement path. The real-life hawk data had many missing steps so we randomly removed 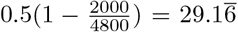 percent of the generated locations. Our simulated paths contained approximately 2000 pairs of consecutive data points (steps), resembling the real-life data. We based the simulated paths on the hawk migration depicted in Figure 2, except we removed the stopover that took place during this migration in some of our simulations. In both stopover and non-stopover cases the simulated birds departed from their breeding grounds after 119 days and arrived on their wintering grounds after 190 days (in line with examples from our results). We simulated 25 paths without any stopover behaviour and 25 paths with a 30-day stopover starting at day 130. We set *ρ*_0_ = 0.6 km, *ρ*_1_ = 7 km, *κ*_0_ = 0, and *κ*_1_ = 0.5. Parameter estimates and information criteria for each simulation can be found in Supplementary File 1.

